# Thermal suppression of gametogenesis can explain historical collapses in larval recruitment

**DOI:** 10.1101/2023.09.28.559919

**Authors:** Daniel K. Okamoto, Nathan B. Spindel, Maya J. Mustermann, Sam Karelitz, Brenna Collicutt, Iria Gimenez, Kate Rolheiser, Evan Cronmiller, Megan Foss, Natalie. Mahara, Dan Swezey, Rachele Ferraro, Laura Rogers-Bennett, Stephen Schroeter

**Affiliations:** Department of Integrative Biology, University of California, Berkeley, Berkeley California, USA 94702; Department of Biological Science, The Florida State University, Tallahassee, Florida 32303; Hakai Institute, Quadra Island, British Columbia, Canada; Coastal Marine Science Institute and Bodega Marine Lab, University of California, Davis California, USA 94923; Kashia Band of Pomo Indians of Stewarts Point Rancheria, Santa Rosa, CA; California Department of Fish and Wildlife; Marine Science Institute, University of California, Santa Barbara

## Abstract

Projections for population viability under climate change are often made using estimates of thermal lethal thresholds. These estimates vary across life history stages and can be valuable for explaining or forecasting shifts in population viability. However, sublethal temperatures can also depress vital rates and shape fluctuations in the reproductive viability of populations. For example, heatwaves may suppress reproduction, leading to recruitment failure before lethal temperatures are reached. Despite a growing awareness of this issue, tying sublethal effects to observed recruitment failure remains a challenge especially in marine environments. We experimentally show that sublethal suppression of female gametogenesis by marine heatwaves can partially explain historical collapses in urchin recruitment. This response differed by sex but was similar between animals from warmer or cooler regions of their range. Overall, we show sublethal thermal sensitivities of reproduction can narrow the thermal envelope for population viability compared to predictions from lethal limits.

## Introduction

Predicting the impact of warming and extreme heatwaves on ecosystems and populations remains one of the most daunting but important tasks in ecology and conservation as climate change impacts intensify. Information about the future dynamics and viability of populations can be gleaned from measuring thermal lethal thresholds (or critical thermal limits - CTLs) in controlled settings, as well as from historical patterns of population performance in nature. For the latter, extrapolating historical patterns without understanding processes that shaped them remains dubious; meanwhile, expectations from CTLs often underpredict the abruptness and magnitude of population responses to warming in nature ^1-3^. For example, warming and heatwaves may lead to recruitment failure in populations even when temperatures remain below lethal thresholds ^4,5^. Thus, the realized thermal niche of many species (i.e., thermal regimes in which a population exhibits viability over time) is often narrower than expected based solely on CTLs for individual life stages (gametes, juveniles, adults, etc. ^6^). Combining inference from a mixture of laboratory studies, historical patterns, and analysis of reproductive rates can provide a more comprehensive understanding of the impacts that climate events such as heatwaves have on recruitment and population dynamics.

Sublethal challenges of climate change and warming are impacting recruitment and subsequent population viability^7^. One pathway by which sublethal temperatures can alter population viability is by affecting gametogenesis, sexual maturation, and/or sterility. Changes in temperature can directly affect reproductive phenology for a diversity of plants and animals^8^ which can significantly impact reproductive success, fitness, and productivity in a warming world. In plants, for example, warmer winters can lead to crop failure by disrupting vernalization pathways. This family of processes, in which plants require specific thermal cues, as well as other environmental patterns or thresholds for induction of flowering and seed/fruit development has long been studied and understood in agricultural and laboratory plant systems (e.g., ^9,10^). Likewise, a diversity of animal taxa can exhibit infertility or low fecundity at sublethal temperatures ^11-15^.

For animals, and marine organisms with planktonic larvae in particular, it remains unclear if and to what extent patterns of recruitment failure in nature are shaped by sublethal thermal suppression of reproduction. This knowledge gap remains for several reasons. First, attributing causality to observed changes in year-class strength remains a challenge for animals with highly dispersive reproductive propagules. Shifts in recruitment may be explained by variation in food supply ^16^, trends in reproductive and mating success ^17^, shifts in dispersal ^18^, or a host of other factors that may also be directly or indirectly related to warming. Second, thermal limits are often estimated using constant temperatures; yet in the oceans, temperature is variable, and the dynamics of this variation can shape patterns of reproduction. Thus, while it is well known that sublethal thermal stress can limit reproduction in marine invertebrates ^19,20^, it remains less well understood if heatwaves can induce such limitations and/or plausibly help explain collapses in recruitment. If climate change and extreme events in nature lead to suppressed gametogenesis well before lethal limits of organisms are reached (e.g., thermal stress that decreases larval, juvenile, or adult survival), then declines in population viability and contractions in distributions may occur and at less extreme temperatures than established under critical thermal limits.

For populations of purple sea urchins (*Strongylocentrotus purpuratus* – a key herbivore) in Southern California, recruitment tends to collapse during warm El Niño events ^21^ (Figure 1A). This phenomenon is remarkably consistent over time (extending back at least six decades), even though temperatures during the reproductive season in Southern California (late fall and early winter ^21,22^ Figure 1B) are almost always below those considered to be lethal for adults (∼25 °C ^23^, < 0.01% of daily observations since 1990 above this value). Moreover, winter temperatures in the Southern California Bight when larvae are present (∼ 11-16°C) are generally well below the thresholds thought to impede multiple phases of larval survival ^24^ (fertilization and early-phase survival of San Diego, Santa Barbara, and British Columbia animals declines to near zero between 20-22°C - SK, unpublished data). Specifically, purple urchins tend to spawn in the winter and spring, coinciding with favorable conditions (low temperatures and high food availability) for larvae ^21^. Larvae persist in the plankton from approximately 28 days to several months depending on food availability and temperature ^25,26^. Adult urchins build gonads over the course of the summer and fall with gametogenesis occurring in the fall and early winter. El Niño conditions in Southern California are characterized by warmer water temperatures in summer, fall, and winter than during neutral years or during La Niña conditions, but still exhibit seasonal cooling (Figure 1B). One proposed explanation for decreases in recruitment in Southern California during El Niño is that temperatures experienced during these events suppress gametogenesis even though energetic investment in gonads may be sufficient for reproduction ^27,28^. Specifically, Pearse ^28^ demonstrated that high temperatures decrease gamete production at sublethal levels (i.e., at 21°C vs 14°C), but the higher temperature exceeded those experienced in the spawning season. Moreover, animals used by Pearse were sourced from cooler regions of the range that rarely experience such temperatures, while animals from southern regions are frequently exposed to temperatures that exceed 21°C.

**Figure 1:**
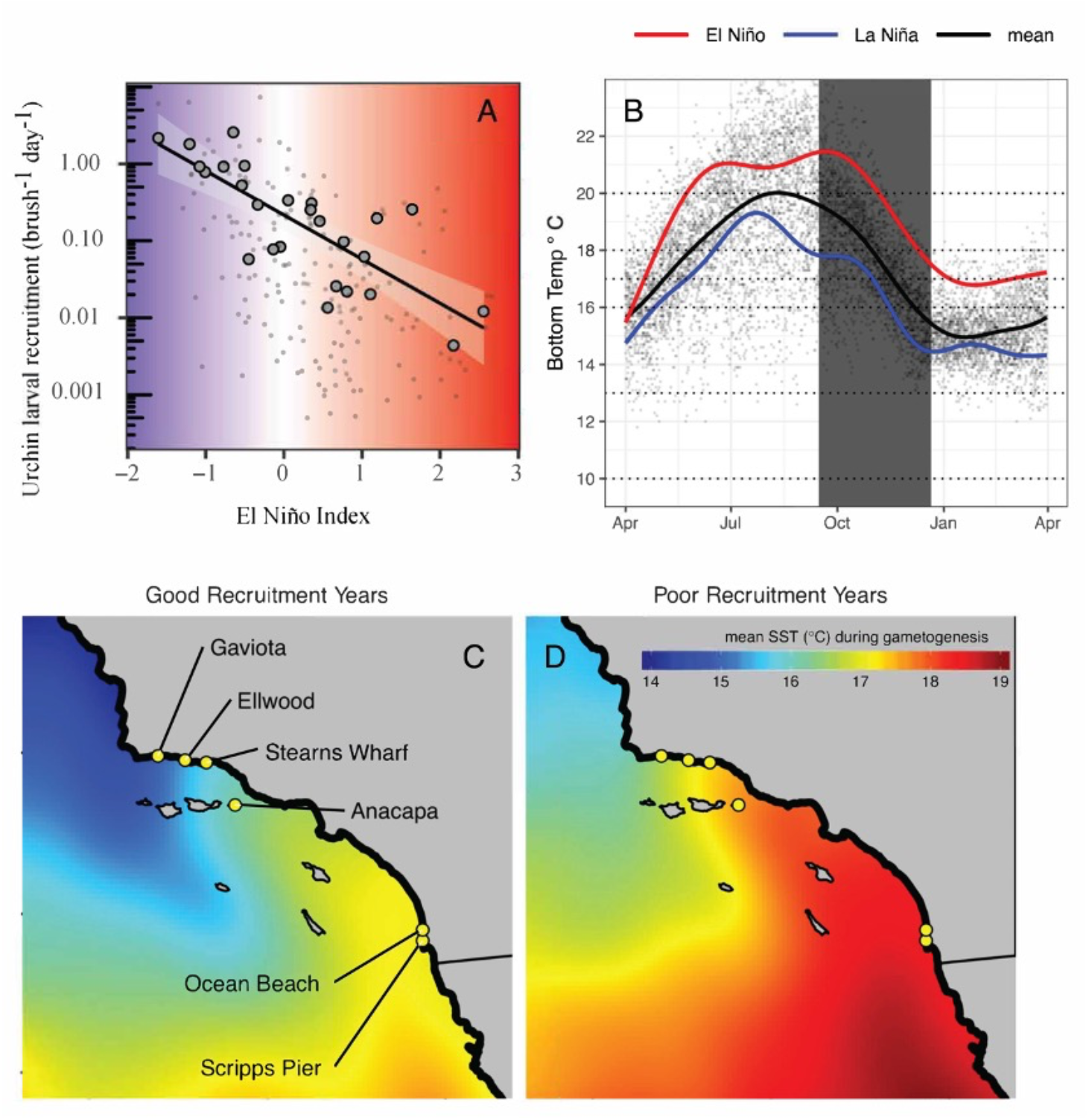
A) Historical mean annual (large points) and monthly mean (small points) larval settlement on standardized brush samplers across six sites in Southern California vs Multivariate El Niño Index (MEI) since 1990 (reproduced from Okamoto et al. ^21^). Colors correspond to the x-axis representing cooler (negative) and warmer (positive) temperatures associated with the MEI in Southern California. B) Historical benthic temperatures from Scripps Pier^53,54^. Small points: daily means. Black line: historical seasonal mean. Red and blue lines: historical El Niño & La Niña seasonal trends, respectively, from the data that were simulated in mesocosms (the period simulated in the experiment is indicated by the vertical grey bar). Horizontal dotted lines: constant mean temperatures simulated in replicate mesocosms. C & D) Mean satellite-derived sea surface temperature from September through December in four strong (C - 1993, 1996, 1999, & 2010) and poor (D - 1994, 1997, 2004, & 2015) years for larval settlement from 1990 to 2016 ^55^. Note that C & D are visualized to show comparative spatial trends, though satellite-derived temperature data may be biased relative to in situ benthic temperature data. Sites in C represent the six historical settlement collection locations that all show negative correlations with the MEI.

**Figure 2:**
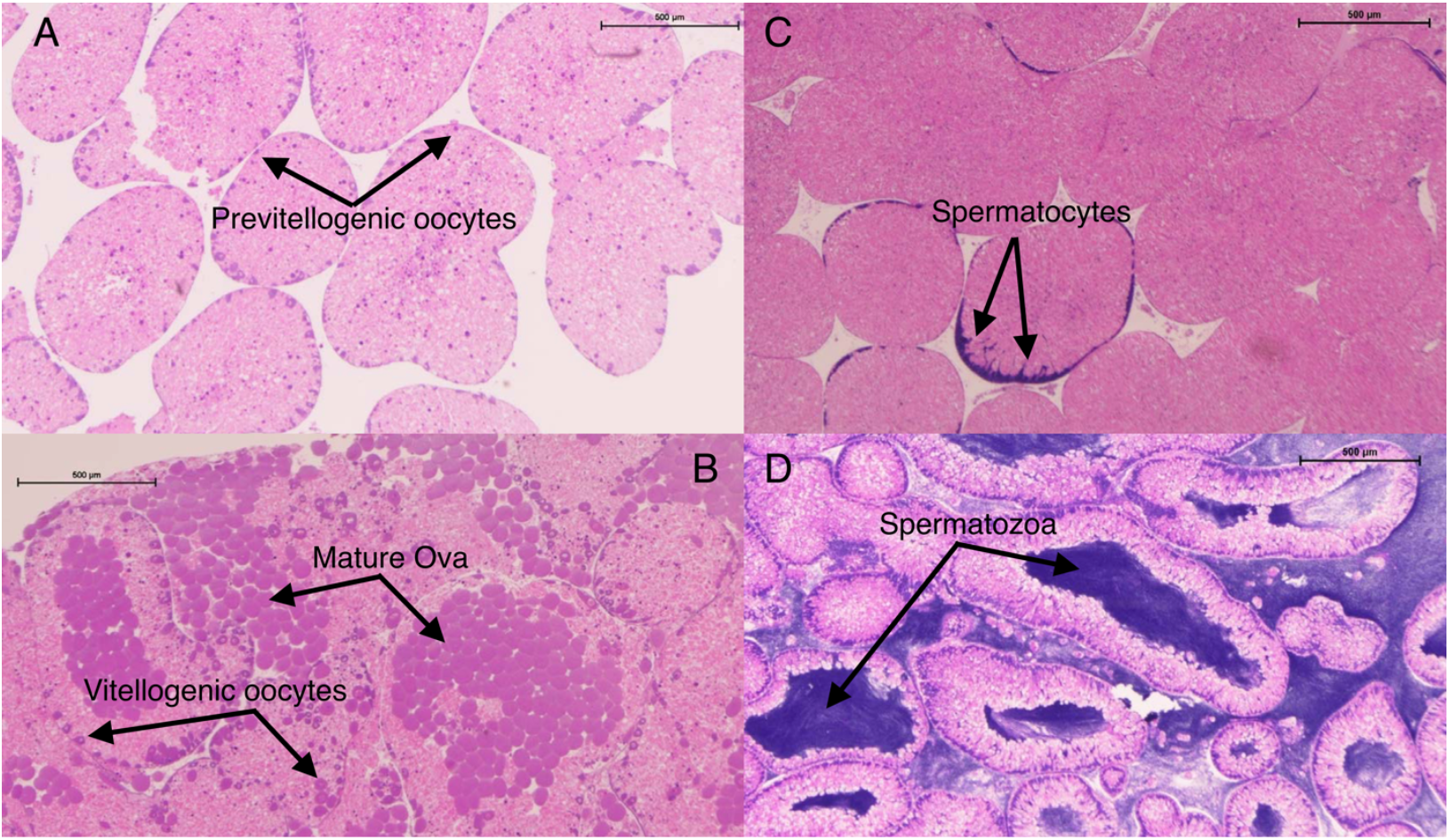
Gonad sections stained with hematoxylin and eosin with scale bar representing 500 μm. A-B Female ovaries in Stage I (A) and Stage IV (B) arrows indicate small previtellogenic oocytes, vitellogenic oocytes in the process of meiosis, and mature ova. C-D. Females in stage I are largely devoid of developing and vitellogenic oocytes with substantial reserves invested in nutritive phagocytes (light pink), Stage II has few developing oocytes, Stage III has some developing oocytes with few, scattered mature eggs, and Stage IV has many mature eggs (dark, solid circles) and some developing oocytes (dark circles with visible, central germinal vesicle). Male testes showing stage I (C) with few developing columns of spermatocytes, and stage IV (D) with large sections of the gonad converted to mature spermatozoa. Males in stage I have few visible developing spermatocytes (columns of dark purple around the margins), males in stage II have dense columns of spermatocytes, males in stage III have few small pockets of spermatozoa, and males in stage IV have large pockets of spermatozoa and rapidly disappearing somatic tissue (light pink).

**Figure 3:**
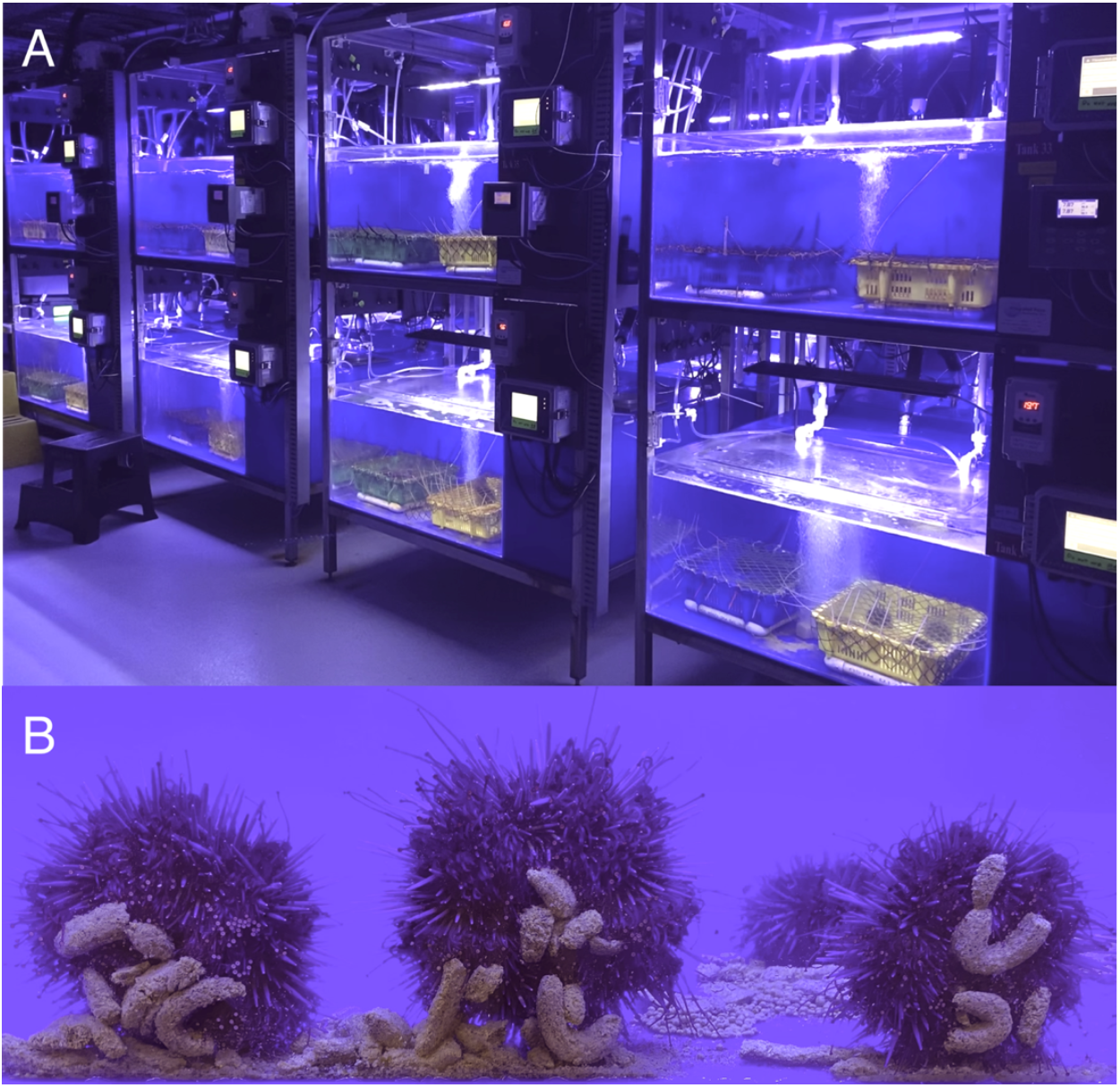
A) Mesocosms at the Hakai Institute’s Quadra Island Ecological Observatory with trays holding experimental subjects. B) Experimental subjects consuming controlled, manufactured Urchinomics algal feed pellets.

In this study, we used *S. purpuratus* to test whether historical collapses in larval supply and year-class strength (i.e., abundance for a given young-of-year cohort) can, in part, be explained by sublethal thermal suppression of reproduction. Specifically, we used two laboratory experiments to ask whether marine heatwaves (prolonged periods of anomalously high temperatures^29^ in this case that correspond with well-defined El Niño is events) that have led to collapses in larval supply also affect reproduction in terms of both gonad production and gametogenesis. For the first experiment we compared effects of simulated heatwaves to simulated historical cool trends, as well as responses under a gradient of constant temperatures. For the second experiment we exposed animals from warm and cool regions to the same treatments in a common garden setting to test whether source population (whether via local adaptation or acclimatization) affected patterns of gametogenesis.

## Results

Simulated El Niño and warmer conditions in the first experiment led to decreased gonad production as well as suppressed gametogenesis in *S. purpuratus*. Declines in gonad production associated with high temperatures were consistent for constant exposure to 20°C and the El Niño treatment that shifted from 21 to 18°C with a mean of 20°C (Figure 4). In contrast, the La Niña treatment (shifting from 18 to 14°C, with a mean of 16°C) led to reduced gonad production compared to a constant mean temperature of 16°C. The effect of treatment on gametogenesis differed depending on whether the thermal environment varied over time (El Niño and La Niña versus constant temperatures) and by sex. With a mean of 16°C, we saw no difference in gametogenesis between La Niña and constant temperatures for males or females. For females, constant high temperature (20°C) and the El Niño treatment (21-18°C with a mean of 20°C) led to a substantial reduction in oogenesis. For males, only the El Niño treatment affected spermatogenesis, with no change observed across constant temperatures from 10 to 20°C. For the second experiment, we found no differences in gametogenesis among source populations in response to treatments, with qualitatively similar treatment effects as in the first experiment for both males and females. Below, we describe the specific results of each experiment in detail.

**Figure 4:**
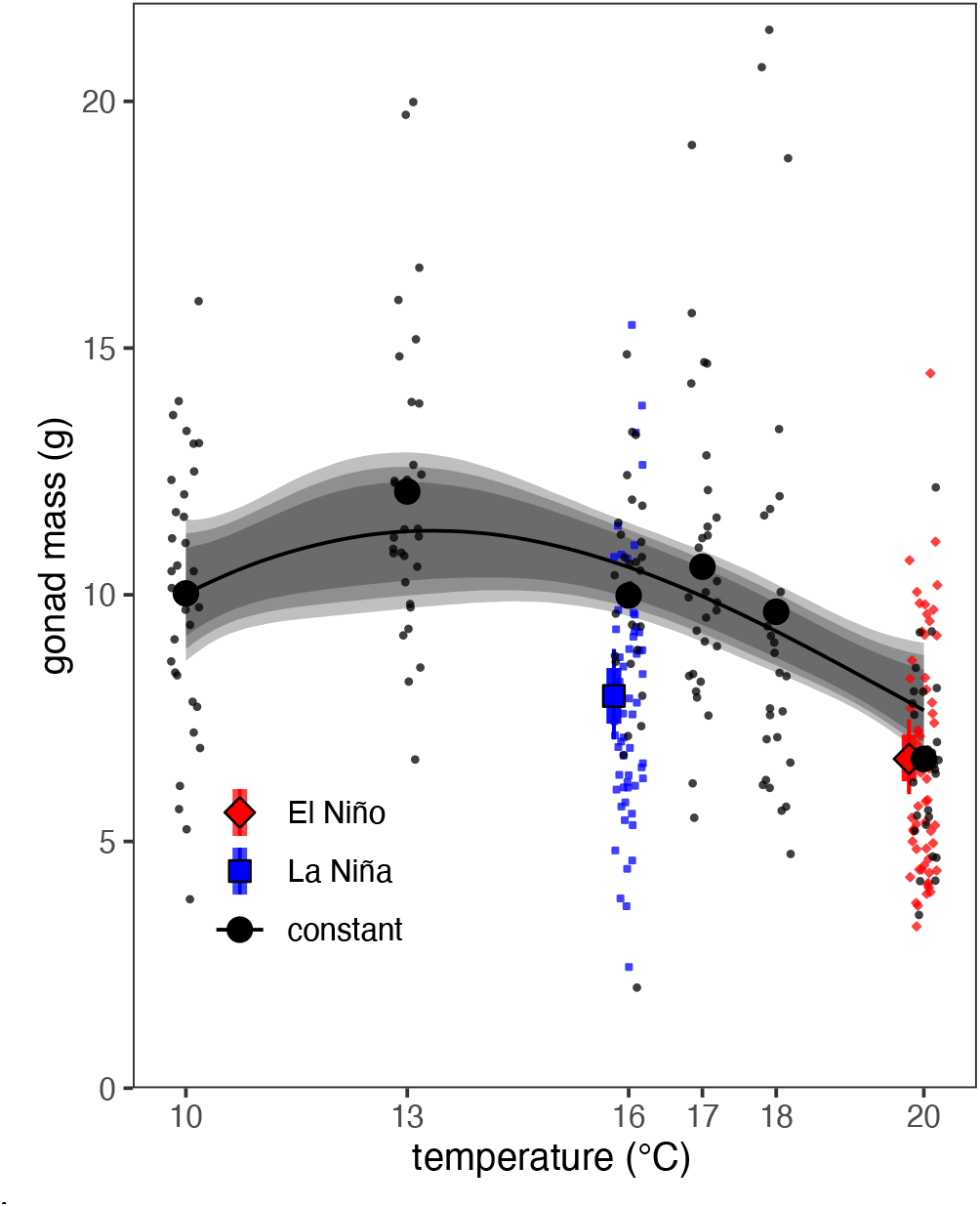
Final gonad mass vs mean temperature. Black circles and the smooth trend represent fixed temperature treatments; the red diamond and blue square represent El Niño and La Niña thermal treatments, respectively. Small points represent individual observations (standardized to 50 mm diameter individual for consistency). Error bars and bands represent the 90, and 95% highest probability density interval of the posterior. Results are shown for males and females combined with no evidence for a difference between sexes in the response (see Results).

### Experiment 1: Effects of El Niño, La Niña, and a thermal gradient on gonad production and gametogenesis

#### Gonad production

In the first experiment, exposure to the highest temperatures (for both variable and constant treatments with mean of 20°C) negatively affected both male and female gonad production (cumulative gonad mass, Figure 4). Specifically, we observed a 42% decrease (95% highest probability density interval [HPD]: 28-55%) in gonad production at 20°C compared to a constant temperature of 13°C (where peak gonad production was observed). Gonad production did not differ for the El Niño (7.03g [95% HPD: 6.20g to 7.96g]) and constant treatment that shared the same mean value of 20°C (7.2g, [95% HPD: 5.97g to 8.50g]). In contrast, the La Niña treatment (declining from 18 to 14°C with mean of 16°C) showed a reduction in gonad mass (26% - HPD: 15 to 38%) compared with a constant value of 16°C (8.69g [95% HPD: 7.68g to 9.77g] vs 11.84g [95% HPD 10.71g to 13.15g], respectively). The lowest temperature (10°C) imposed a smaller, but still significant reduction in gonad mass (22% reduction, 95% HPD: 4-39%) compared to the peak at 13°C. The reduction in gonad production from the La Niña to El Niño (with means of 16°C and 20°C, respectively) was smaller than when comparing constant temperature treatments of 16°C to 20°C (17% [HPD: 3.5 to 32%] vs 38% [HPD: 23 to 39%] respectively) despite having the same mean temperatures. This reduced effect resulted from the aforementioned decline in gonad size in La Niña versus equivalent but constant mean of 16°C. Notably, we saw no difference in gonad production by sex. The final model used for analysis (temperature treatment but no sex or sex x treatment interaction) is supported by Bayes Factors >100 for temperature treatment relative to both the null and to models with sex and interactions. Bayes Factors > 100 are considered decisive evidence of one model over a second, values 10-100 are considered strong evidence ^30^, while values <1 favor the second model.

#### Gametogenesis

The El Niño treatment led to a significant reduction in gametogenesis for both males and females, and this effect was more extreme for females compared to males. We scored gametogenesis using a standard staging scale of I to IV where IV is fully mature and I is immature (sensu ^31^). Specifically, far fewer individuals in the El Niño treatment reached Stage IV (most mature) compared to the La Niña treatment for both males and females (Figure 5). Yet this effect was more extreme for females compared to males. Female urchins incubated in the La Niña treatment actively produced mature gametes (89% [95% HPD: 77 to 94%] of females in stage IV, Figure 5A). In contrast, only 15% [95% HPD: 4 to. 28%] of El Niño females were in stage IV (Figure 5A), while most females were in stage I. This decline in gametogenesis associated with El Niño amounted to an estimated 74% reduction for females compared to La Niña (Figure 5A, HPD: 56 to 91% decline in females in stage IV). For male urchins, these results were consistent but less extreme. 92% of La Niña males were stage IV [HPD: 77 to 99%] vs 44% of El Niño females [HPD: 23 to 67%]– a reduction of 48% [HPD: 23 to 72%] - Figure 5B). Overall, models that included the treatment, sex, and interaction compared to the models with only sex were supported Bayes Factors >100.

**Figure 5:**
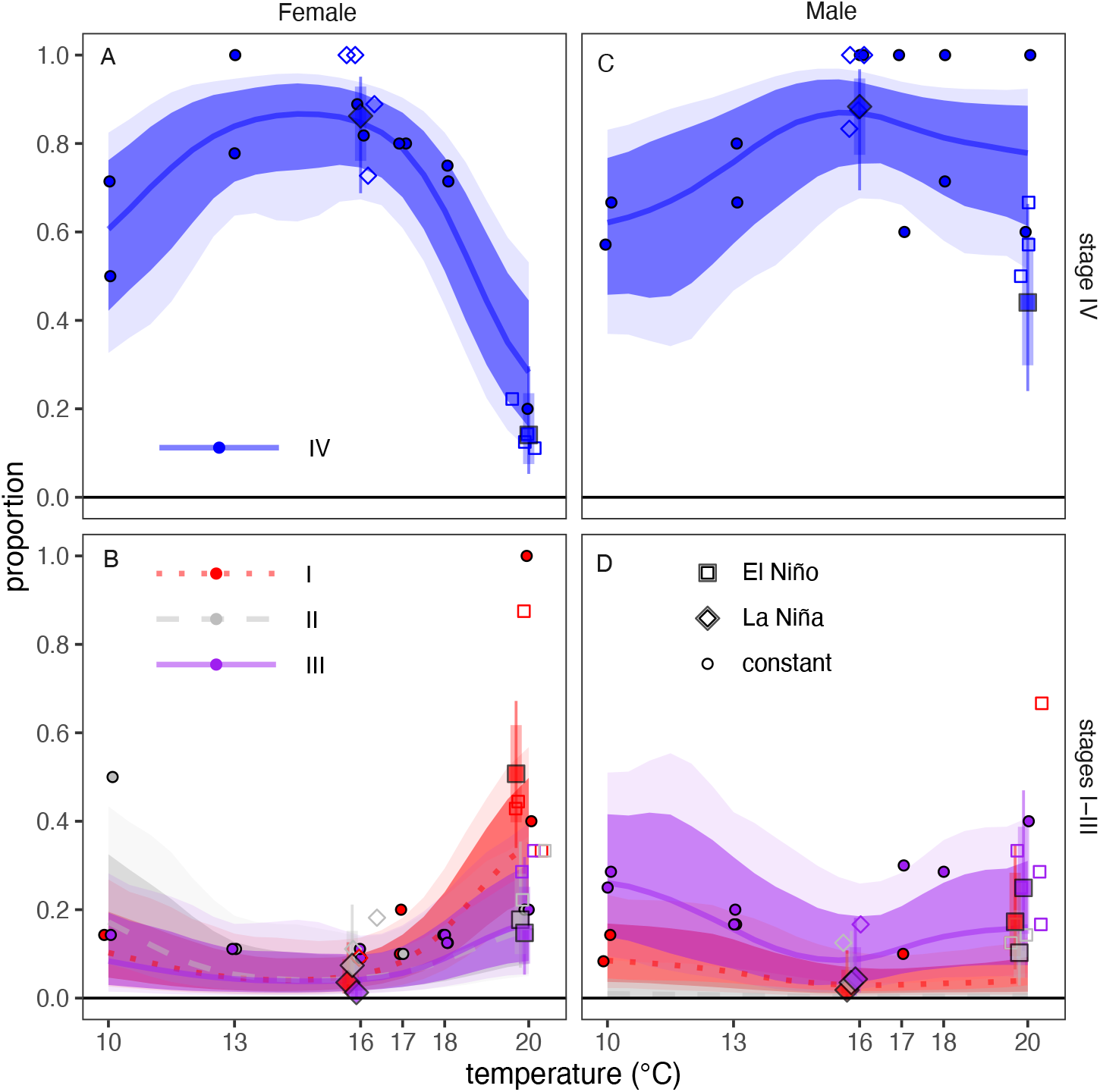
Proportion of adult females (A, B) and males (C, D) in each reproductive stage by treatment (El Niño (squares), La Niña (diamonds), or constant temperature (circles)) and mean temperature. A) Proportion of females in stage IV. B) Proportion of females in stages I (dotted red lines, red points), II (dashed grey lines, grey points, and III (solid purple lines, purple points). C) Proportion of males in stage IV, and D) proportion of males in stages I, II, and III (same symbology as females). Circles represent empirical means for constant temperature treatments, while El Niño and La Niña points represent posterior mean estimates. Constant temperature treatments had two replicate mesocosms per treatment and 15 animals per mesocosm. El Niño and La Niña had four replicate mesocosms and 15 animals per mesocosm. Uncertainty bands/intervals represent 95% (light/thin) and 90% (dark/thick) from the ordinal logistic regression.

Our results from the first experiment also indicate that temperature dynamics, and not just their mean values, can alter reproductive response depending on sex. Simulating a historical El Niño trend resulted in a reduction in the proportion of males investing in spermatogenesis in contrast to no change in spermatogenesis under a constant mean temperature. Specifically, male animals subjected to the El Niño treatment (simulated seasonal peak and decrease from 21 to 18°C with a mean of 20°C) compared to a constant 20°C throughout showed substantially lower rates of full gonad maturation (stage IV) with a reduction from 81.2 to 47.8% (a difference of 37% [95% HPD: 8% to 65%], Figure 5C, D). In contrast, we observed similarly low rates of full maturity for females subjected to the El Niño simulation and constant 20°C treatment (15% for El Niño vs 25% for constant 20°C, Bayesian P = 0.28, (Figure 5A, B).

### Experiment 2: Effects of source population on effects of El Niño on gametogenesis

When comparing populations, we found a similar strong effect of temperature treatment on gametogenesis but did not find a significant difference among populations in these responses (Figure 6). Specifically, we did not find differences in gametogenesis in response to 20°C or El Niño conditions (relative to 10°C) among animals sourced from different populations, including those from warm regions (San Diego County and Santa Barbara County) or a cold upwelling region (Sonoma County). Comparing models using Bayes Factors found no support for the hypothesis that treatment effects varied by source population (Bayes Factor = 19 favoring the reduced model [treatment, sex, and interaction] compared to the full model [with location, treatment, sex and interactions]). The lack of location of origin effect is reinforced by analysis of the full model, where we found no difference among locations in estimated treatment effects for any stage (i.e., 95% credible intervals overlap zero for differences among populations in the log-scale treatment effects for either 20°C or El Niño relative to 10°C by sex); however, we did find consistent effects of each treatment across locations (>95% posterior probability of a difference in proportion of stage IV animals for both 20°C and El Niño relative to 10°C for females and El Niño relative to 10°C only for males). Thus, hereafter we describe analysis using the model without location interactions. Overall, we estimate a combined 75% and 90% reduction in stage IV females and males (respectively) for the El Niño treatment relative to the 10°C treatment (95% HPD = 92% to 61% reduction for females and 99% to 65% reduction for males) and an 81% and 96% reduction in stage for IV females and males for the 20°C treatment relative to the 10 degree treatment (95% HPD = 96% to 68% reduction for females and 99 to 72% reduction for males). The pooled results are shown as the thick lines/points in Figure 6 D and H for females and males, respectively. While results from the first experiment were conducted with animals sourced from cooler water, gametogenesis trends are similar when comparing animals from warmer versus cooler environments.

**Figure 6:**
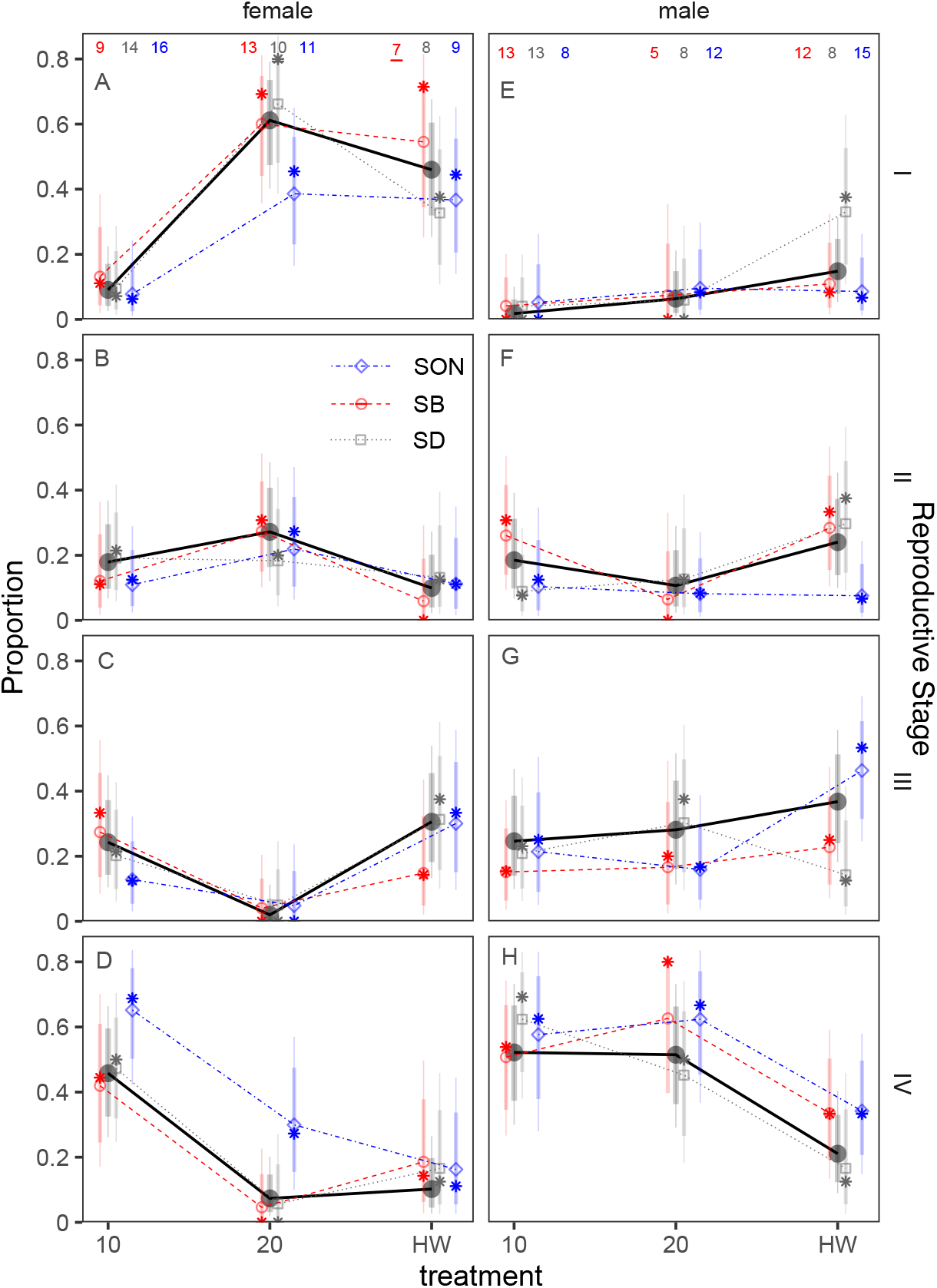
Estimated proportions of each stage (rows) for each sex (columns) for each treatment from the population comparison experiment (Sonoma, Santa Barbara, San Diego). Points represent mean and error bars 90/95% credible intervals. The model with no location interaction is indicated by black circles while other symbols/colors reflect the model with location interaction with sex/treatment (red circles - SB = Santa Barbara County, grey squares - SD = San Diego, and blue diamonds - SON = Sonoma). All estimates are derived from Bayesian ordinal logistic regressions with random effects of location and vague priors on the estimated means. Sample sizes are shown in panels A and E for males and females, respectively. Asterisks represent empirical mean proportions.

## Discussion

Global warming and extreme heatwaves present a significant threat to ecosystems worldwide. Characterizing temperature thresholds that cause mortality across life stages can provide essential insight into the impacts of heatwaves. It is well understood that range limits can contract and abundances can collapse well before organismal lethal limits are reached. Mechanistically explaining how and why such sublethal thermal regimes affect populations remains a challenge. In this study, we show how thermal suppression of reproduction can, in part, provide a plausible explanation for historical collapses in larval supply associated with warming events. Our results provide two key general conclusions. First, sublethal impacts of warming events on reproduction may lead to recruitment failure and population declines before lethal upper thermal limits are experienced. This insight is derived from the results of our lab experiments in the context of our historical observations of larval supply. We found these results to be consistent for animals sourced from both warmer and cooler parts of their range. Second, variable temperature treatments that reflect historical observations of heatwaves, in comparison to constant temperatures, exert a more pronounced effect on reproductive suppression. Therefore, ignoring the dynamics of thermal regimes may underpredict the magnitude of sublethal thermal effects. Importantly, these results raise questions regarding physiological processes by which temperature shapes gametogenesis, and how such effects may vary in and among populations over time. These effects will almost certainly interact with other stressors such as food availability for adults (sensu ^32^) or shifts in physical dispersal, gametogenesis, or heatwave conditions.

Organismal thermal lethal limits can underpredict observed shifts in population-level vital rates. Our experimental results provide plausible evidence that sublethal reproductive failure contributes to collapses in year-class strength. These insights hinge upon a combination of long-term monitoring, historical observational data collection, and incremental prior experimentation. Specifically, our historical data collection in Southern California shows larval supply of purple urchins consistently collapses during marine heatwaves^21^. Prior field observations proposed that temperatures above a 17°C threshold led to suppressed reproduction during a marine heatwave (assessed observationally via seasonal gamete extrusion ^33^ and histology ^27,33^). Additionally, the prior experiment by Pearse ^28^ in Monterey Bay, California showed that urchins exposed to constant 21°C suppressed male and female gametogenesis compared to constant 14°C. However, 21°C is outside the range of temperatures generally experienced during the spawning season, even during marine heatwaves in Southern California (Figure 1A, Figure 7A,B). Our results suggest a higher limit than 17°C but still support the notion that El Niño can suppress gametogenesis. Specifically, replicating historical marine heatwaves (shifting from 21°C and ending at 18°C with a mean of 20°C) can suppress gametogenesis for both males and females sourced from across a wide latitudinal range. For females, this suppression appears to occur above 18°C but at or below 20°C. Females exposed to a constant 18°C in the first experiment proceeded with vitellogenesis and produced mature gametes, whereas females generally did not when under the El Niño treatment (ending at 18°C) or the constant 20°C treatment. Males exposed to a constant 20°C proceeded with spermatogenesis and produced mature gametes but not those in the El Niño treatments. In contrast to gametogenesis, overall treatment effects on gonad production (mass – a metric of potential fecundity) did not differ by sex. These disparities in gametogenesis patterns among sexes raise several hypotheses that require further investigation. For example, sperm and eggs may differ in their thermal constraints, time required to complete gametogenesis, and/or costs associated with production and maturation. Despite the aforementioned differences, El Niño treatments that ended at 18°C yielded consistent results for gametogenesis among males and females. This final temperature is several degrees lower than what is considered to be lethal for adults and larvae (23-25°C and 20-22°C, respectively). As a result, inferences derived solely from larval or adult survival under laboratory conditions would not predict collapses in year-class strength in association with marine heatwaves.

**Figure 7:**
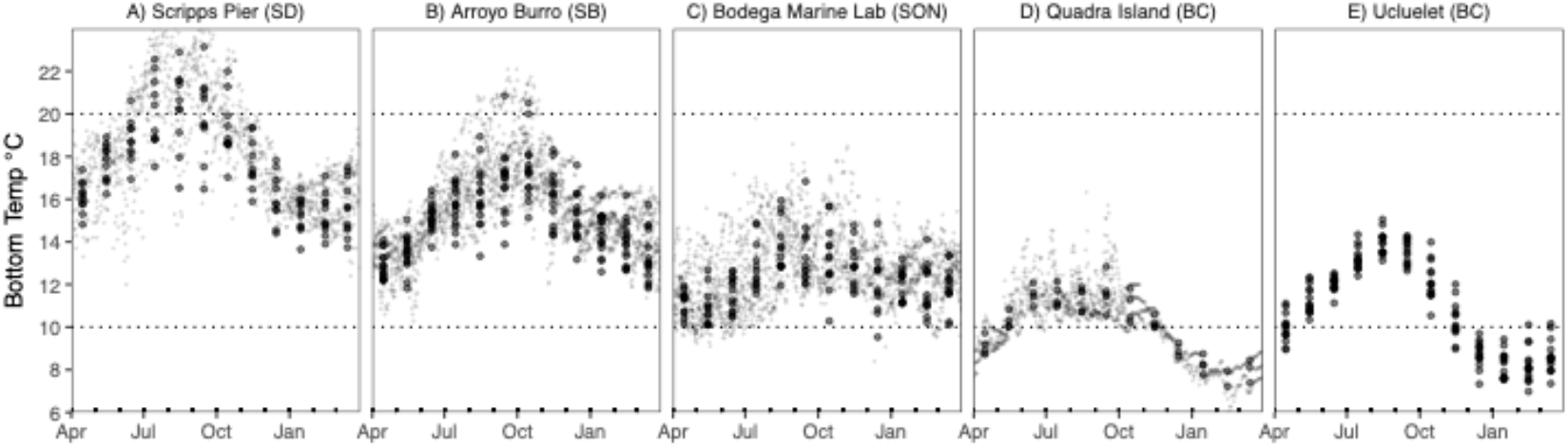
Daily benthic temperatures from San Diego (Scripps Pier), Santa Barbara (Arroyo Burro SBC LTER), Sonoma (Bodega Marine Lab), D) the Quadra Island Ecological Observatory, BC (the site of the Marna experimental work) and E) surface temperatures for the Amphitrite Point lighthouse in Ucluelet, BC (the source of the animals for the Marna experimental work). Small points represent daily means (except in E where original data are monthly values) and large points represent monthly means. For consistency, data are shown only since 2015 (the inception of the Quadra Island time series).

Heatwave events in both marine and terrestrial settings are often highly dynamic, characterized by rapid fluctuations in temperature. We’ve shown how experiments that test constant temperatures and ignore such fluctuations may yield erroneous conclusions. When comparing variable versus constant temperature treatments, gonad production (and by extension potential male and female fecundity) differed between La Niña and constant temperatures of the same mean. These disparities in gonad production likely result from nonlinear averaging (e.g., Jensen’s inequality) associated with dynamics of feeding, metabolism, and/or structural growth allocation ^34,35^. With regards to gametogenesis, male but not female patterns of gametogenesis consistently differed between constant versus El Niño temperature profiles that shared the same mean value (20°C). Thus, the variability, specific time course, and extremes of environmental exposure have the potential to alter reproductive responses to heatwaves and can differ by sex. Importantly, our studies ended in December, when temperatures approach winter lows (Figure 1, Figure 7). Even in El Niño years, winter temperatures are likely sufficient to facilitate reproduction over the long term (Figure 1). Thus, rather than inducing skip-spawning, thermal suppression of gametogenesis in fall and early winter may instead lead to a shortened spawning season. Because urchin settlement timing aligns with peak phytoplankton production^21^, such effects may lead to other negative impacts on recruitment such as match-mismatch dynamics with poor survival ^36^. Moreover, our variable treatments are limited to a specific set of trajectories and our constant treatments do not include 21°C or above (the maximum temperature of the El Niño treatment); thus, future work may benefit from assessing how brief or consistent incursions into the 21°C range, and the timing of such incursions, affect both male spermatogenesis and female oogenesis. As a result, future studies may benefit from documenting gonad production and gametogenesis under different real-world and forecasted conditions in both controlled settings and among populations across time and space.

Many species exhibit spatial and/or population level variation in response to temperature. Purple urchins, for example, can display significant differences in environmental sensitivities among populations (e.g., ^37^). Our results suggest that, at least for the populations sampled and treatments applied, impacts of marine heatwaves are consistent between sea urchins sourced from cool and warm regions of the range. These results reflect only these specific contrasts and populations, and thus cannot rule out variation to different contrasts or variation in the response in other populations. Yet with animals ranging from British Columbia to Southern California, we show that marine heatwave effects on sea urchin gametogenesis are likely significant in those regions exposed to warm waters. However, we acknowledge that linking these results to outcomes in the field definitively, rather than logically, requires documenting the spatial and temporal patterns of reproductive phenology and gametogenesis in nature and any association with larval supply. Moreover, the physiological and regulatory processes shaping these patterns have yet to be systematically characterized. Thus, examining the genetic and plastic basis of how reproduction and gametogenic cycles may respond to changes in temperature regimes remains an important objective both in this species and more broadly.

These experiments present plausible evidence for how and why historical collapses in larval supply occur during marine heatwaves. Yet there are likely myriad factors that occur during marine heatwaves that may shape recruitment in addition to, or in synergy with suppression of reproduction. First, food stress occurs during El Niño events in Southern California ^38^. Such trends might exacerbate the effects of temperature on gamete production as food limitation for adults may alter thermal energetic performance or allocation (e.g., via “metabolic meltdown” ^32^). In these experiments, we focused on well-fed adult animals to avoid confounding effects of food availability. The results shown here raise the question of whether declines in availability and productivity of macroalgae can exacerbate responses of gametogenesis in sea urchins. Second, El Niño events are thought to be coupled with shifts in ocean circulation that might alter patterns of phytoplankton production (i.e., larval food) as well as the delivery of planktonic larvae to suitable settlement sites. These trends may also affect reproduction as urchin spawning may be sensitive to changes in phytoplankton-derived chlorophyll ^39,40^. Thus, even in the cases where offspring are produced from successful maturation and spawning, shifts in larval survival and delivery may further compound effects of heatwaves. Finally, temperature ^41^ and animal density may increase disease prevalence and disease related mortality ^42,43^ in nature in addition to the reproductive effects observed here. Therefore, our results focus on one set of key factors among many that may simultaneously respond to abiotic and biotic conditions during extreme events known to alter dynamics of populations.

Marine heatwaves have led to substantial reorganization of ecosystems. This trend has been particularly apparent in recent years in temperate rocky reefs ^44-50^ with resulting, negative effects on ecosystem services and biodiversity ^51^. While extreme events can impose direct and observable effects on mortality, sublethal effects, when present, may be far more frequent and insidious because they occur below lethal thresholds and may have less immediately visible or observable impacts. This suggests that survivors of marine heatwaves may be impaired and reproductively dysfunctional despite being alive ^19^. This study is among the first to show that widespread, historical collapses in sea urchin larval supply in the field can at least partially be explained by sublethal suppression of gametogenesis resulting from marine heatwaves. Moreover, our study demonstrates the value of long-term population studies and quantifying non-lethal physiological and reproductive sensitivities when considering sublethal thermal envelopes or thermal fertility limits ^52^ and the population-level impacts of global warming and heatwaves.

## Methods

To quantify how different thermal regimes affect investment in gonads and development of gametes in male and female urchins, we first conducted a 10-week experiment in which 300 animals were incubated in replicate 350L mesocosms that simulated El Niño (N = 4 mesocosms, 60 animals per treatment) or La Niña (N = 4 mesocosms, 60 animals) conditions based on historical, empirical benthic temperature time series from Scripps Pier in La Jolla, California ^53,54^(trends: Figure 1B, map: Figure 1C) that coincide with historical collapses in larval supply in Southern California. We paired these treatments with a range of fixed temperature incubations (10, 13, 16, 17, 18, 20 ^°^C, N = 2 mesocosms, 30 animals per treatment), two of which matched the mean temperature of the El Niño (20 ^°^C) and La Niña (16 ^°^C) (Figure 1). We chose this benthic time series rather than satellite-derived sea surface temperature information because sea surface temperatures can be markedly different than temperatures experienced at depth by benthic organisms^55^. Experiments were conducted at the Marna Lab at the Hakai Institute’s Quadra Island Ecological Observatory in Heriot Bay, British Columbia due to availability of sophisticated seawater systems for precisely controlled and replicated temperature manipulations. However, animals in this first experiment were sourced from populations found in Ucluelet, British Columbia that rarely if ever experience the high temperatures commonly observed in southern California. Thus, we also conducted a second experiment in which animals were sourced from warm and cool populations and compared responses of urchins to different thermal treatments.

To quantify the degree to which animal source location affected results, we conducted a second 12-week experiment comparing animals from the warmer Southern California bight including San Diego and Santa Barbara with the cool upwelling region of Sonoma County. Seasonal daily temperature data from 2009 (the earliest record at Bodega Marine Lab) for all sites are shown in Figure 7. Seasonal averages from September into December since 2009 are 19^°^C, 16.8 ^°^C, and 13.5 ^°^C from Scripps Pier (San Diego), Arroyo Burro (Santa Barbara), and Bodega Marine Lab (Sonoma), respectively. The percent of daily observations above 19^°^C at these sites since 1990 are 47%, 10%, and <1% at Scripps Pier, Arroyo Burro, and Sonoma, respectively. In this experiment, we exposed animals to the same heatwave treatments (21-18^°^C), 10^°^C, and 20^°^C in a split plot design. See below for experimental details.

For both experiments we used the same general methodologies (food source, feeding regime, containers) but in different settings (Quadra Island Ecological Observatory in experiment 1 and Bodega Marine Lab in experiment 2).

### Field Collections and Acclimation

For the first experiment, we collected sea urchins by hand on SCUBA in the vicinity of Ucluelet, British Columbia, Canada (48.94°N, 125.56° W) from a depth of 7-8 m relative to mean low tide in September 2021 and transported them to the Marna Lab via truck in seawater filled coolers with bubblers in less than 24 hours. We transferred sea urchins to flow-through sea tables and allowed them to recover for a period of one week before placing animals into the mesocosm system. We selected healthy individuals within a constrained size range for incubations (n = 300, mean test diameter = 56.09 mm, range test diameter = 42.12 – 69.46 mm). Finally, we assigned animals to mesocosms at random at ambient temperature and exposed each assigned mesocosm to a temperature ramp, where the ramp reached target temperatures after two weeks from the initial incoming, ambient temperature (mean across all tanks of 13.3°C, SD = 0.3°C) to avoid thermal shock. Once initial target temperatures were reached, they were maintained or, for the variable treatments, were manually adjusted daily in the AM (∼8am each day) as needed by 0.5 ^°^C increments in a scheduled manner to match historical mean El Niño and La Niña daily temperature trends shown in <u>Figure 1</u>B.

For the second experiment, we collected animals from three locations in California including Stillwater Cove in Sonoma County, Mohawk Reef in Santa Barbara, and Point Loma in San Diego using SCUBA from 3-5 meters mean low water. Urchins were dry transported layered between kelp and transferred via ground transportation and into the ambient flow through sea water tanks at Bodega Marine Lab within 24 hours of collection. Temperature ramps and acclimation periods were manually adjusted in 1ºC increments.

### Mesocosm Systems

For the first experiment, we placed urchins in a custom-built array of twenty replicated 214 L [90(L) x 59.5(W) x 40(H) cm] acrylic mesocosms supplied with flow-through UV sterilized and filtered seawater (Figure 3). Each mesocosm was capable of independent control of temperature and animals were provided a lighting regime for all mesocosms using LED fixtures (Aquamaxx, CA, USA) programmed to provide 10L:14D with two-hour linear light intensity transition periods for dawn and dusk (0-100% from 07:00 to 09:00 “dawn”, and 100-0% from 17:00 to 19:00 “dusk”). Each mesocosm independently maintained temperature treatments using a heat exchanger fitted with a titanium coil regulated by a dual stage digital temperature controller (Resolution = 0.1°C, Dwyer Instruments, LLC.^©^, Michigan City, IN, USA). The mesocosm system employed central cooling (Aermec Mits Airconditioning Inc., Mississauga, ON, Canada) and heating (boiler array, Viessmann Manufacturing Company Inc., Warwick, RI, USA) to supply independent heat exchangers with on-demand cold and warm glycol loops for down- and up-regulation of water temperature, respectively. We manually checked and re-calibrated sensors, as needed, using digital traceable thermometers twice daily to control potential temperature sensor drift. We randomly assigned mesocosms to the specified treatments.

For the second experiment, we placed individuals in a custom-built experimental array at the Bodega Marine Laboratory in which individuals from each population were placed in a split plot design in replicated (N = 4 each 140L - 73 (W) x 66 (H) x 32 (D) cm) acrylic tanks per treatment (Oceans Design Inc). Tanks were fed by sumps fixed with both 0.25hp chillers (Aqualogic Delta Star®) and 1000W heat sticks (TSHTCE-1000S) and temperature controllers to regulate temperature in a partially recirculating flow through system with fresh seawater allocated to each system at a rate of 0.5L/min (approx. 5x turnover per day). Each sump was affixed with protein skimmers, UV filter, and bio-ball filters such that water was first filtered and then UV sterilized upon recirculation. Temperatures were set by hand each day for the heatwave treatments and checked with temperature probes for all treatments. We ran experiments from September 15, 2023, to December 19, 2023.

### Animal husbandry

For both experiments, animals were fed on the same feeding regime and the same food as above, also housed in the same density and same trays for consistency. We fed individuals uniform dry pellets combining several macroalgal species formulated for the aquaculture of *S. purpuratus* (Urchinomics Canada Inc., Halifax, NS, Canada). Animals in mesocosms were fed twice per week and we removed uneaten food and refuse every 72 h. Food rations were determined by trial and error in a prior pilot study to ensure all animals had consistent access to food over time. To optimize access for all mesocosm inhabitants to abundant food, we enclosed subjects and food in aquaculture baskets (two baskets per mesocosm, 7 or 8 animals per basket, Thunderbird Plastics 48 x 33.5 x 10 cm Fish Farm Tray) such that food was always readily accessible, and movement was not impeded. Each animal was supplied approximately 2.7 grams of pelleted food twice per week (either 19 or 21 grams per basket for the baskets with 7 and 8 individuals, respectively in each mesocosm) for the duration of the experiment.

### Histological and gonad assays

At the end of the first experiment, we measured all individuals to test for changes in height and diameter (using precision digital calipers) and wet mass (to the nearest 0.1 g). Full growth measurements are reported by Spindel et al. (2023)^34^. Animals were then sacrificed to measure gonad and histological properties. After opening urchin tests, we immediately removed gonads for sampling. We excised one gonad from each animal for histological analyses; a second gonad was excised and carefully weighed to the nearest 0.01 g. Using a clean, sterile scalpel we excised an approximately 2 mm cross section from the first gonad which we immediately placed in a histological cassette, preserved in Hartmann’s fixative for 24 hours, and transferred to 70% EtOH. Preserved gonads were embedded in paraffin, sliced, stained using eosin and hematoxylin, and mounted. We assessed gonad samples for sex and developmental stage using four visual subsections and the entire sample collectively to ensure agreement among subsamples. Histological slides were scored on a scale of I to IV (sensu ^31^), where representative stages are depicted in Figure 2. For the second experiment, we only report analyses of histological data in this paper.

### Statistics and Reproducibility

For the first experiment, we estimated the probability of individuals in each reproductive stage as a function of mean temperature by sex and in response to El Niño and La Niña treatments using a Bayesian multinomial logistic regression model. We modeled the response to mean temperature using a non-parametric Gaussian process smoother. To account for treatment effects, we included a categorical effect of treatment (constant, El Niño, La Niña). Gonad mass was modeled with a Gamma likelihood and log link with the log of test diameter included as a covariate to account for gonad-body size allometry. Because gonad mass was expected to potentially exhibit a concave response to temperature (i.e., consistent with a constrained thermal performance curve), a 3^rd^ order polynomial was used for mean temperature. For both analyses, intercepts were allowed to vary randomly by mesocosm. For the second experiment, we used the same model structure for reproductive stage but with no smoother effects (only categorical terms for constant 10°C, constant 20°C, or El Niño) and included location of collection as a random effect.

For both datasets, we directly compared parameter posteriors within models and to ensure that comparing parameters within models was robust to model (structural) uncertainty, we conducted model comparison using Bayes Factors^56^ (as hypothesis tests of the full versus null models). For Bayes Factors we estimated the marginal likelihood using bridge sampling^57^ via the R package bridgesampling^30^. Because Bayes Factors can be sensitive to prior construction, we ensured our analyses are robust by simulating 200 datasets from both the full and null model under the same prior construction to ensure that Bayes Factor would consistently reject the null when simulating from the full model and vice versa. We sampled posteriors in Stan^58^ (via brms^59^ for Gamma regressions for gonads or coded directly for multinomial models of histology) in R. We ran models with vague priors, and conducted sensitivity analyses to ensure varying prior distributions and hyperparameters did not qualitatively affect the results. For the multinomial models, we provided stage proportions by treatment for the multinomial models vague Dirichlet priors (all concentration parameters = 1 yielding equal prior odds with high uncertainty).

Location random effects were provided logit-scale normal effects (with mean zero) from the mean stage proportions where the scale hyperparameter was provided a folded student-t (df = 3, lower bound = 0, scale = 0.1) hyperprior. Finally, for the Marna Lab dataset, the logit-scale Gaussian process model was given a squared exponential covariance and with length and scale hyperparameters having folded normal hyperpriors (both with sd = 1 and mean and lower bound of 1 and 0, respectively). We ran models for 12000 iterations for each of four chains after a 2000 iteration warmup period and checked convergence visually, that parameters had convergence diagnostics (Rhat) less than 1.001. Data and metadata from this study are available at https://www.bco-dmo.org/project/818918 and analysis and code is available at https://doi.org/10.5281/zenodo.15420518.

## Author contributions

DKO, NBS, MJM, DS, BC, LRB and SS designed the research. DKO, NBS, MJM, BC, SK, KR, DS, IG, EC, MF, and NM conducted mesocosm experiments. RF, MM, and DKO conducted histological scoring. DKO conducted statistical analyses. DKO wrote the initial draft of the manuscript, and all authors contributed to revisions.

## Acknowledgements

This study was funded with generous support from the Hakai Institute, a program of the Tula Foundation, a grant from the US National Science Foundation to DKO and LRB (OCE nos. 2023649 and 2023664), as well as from the Santa Barbara Coastal (SCB) LTER (NSF OCE no.1831937). We thank Eric Peterson, Christina Munck, as well as the facilities staff at the Quadra Island Ecological Observatory for their generous support. Ellington Chen, Janelle Beduya, and Rebecca Ward-Diorio assisted with imaging histological slides. Histology Consultants Inc. conducted histological sectioning. We thank the Coastal Marine Science Inst. and Bodega Marine Lab at UC Davis as well as the California Dept. Fish and Wildlife LRB. Rachel Simons, Wiley Evans, and two anonymous reviewers provided useful feedback on prior versions of the paper.

## References

1 Braschler, B., Chown, S. L. & Duffy, G. A. Sub-critical limits are viable alternatives to critical thermal limits. J Therm Biol 101, 103106 (2021). 10.1016/j.jtherbio.2021.103106 PMID - 34879920

2 Kinzner, M. T. et al. Is temperature preference in the laboratory ecologically relevant for the field? The case of Drosophila nigrosparsa. Global Ecol Conservation 18, e00638 (2019). 10.1016/j.gecco.2019.e00638

3 Rezende, E. L., Bozinovic, F., Szilágyi, A. & Santos, M. Predicting temperature mortality and selection in natural Drosophila populations. Science 369, 1242–1245 (2020). 10.1126/science.aba9287 PMID - 32883867

4 Oliver, E. C. et al. Marine heatwaves off eastern Tasmania: Trends, interannual variability, and predictability. Progress in Oceanography 161, 116–130 (2018).

5 Munday, P. L., Jones, G. P., Pratchett, M. S. & Williams, A. J. Climate change and the future for coral reef fishes. Fish and Fisheries 9, 261–285 (2008).

6 Terblanche, J. S., Deere, J. A., Clusella-Trullas, S., Janion, C. & Chown, S. L. Critical thermal limits depend on methodological context. Proceedings of the Royal Society B: Biological Sciences 274, 2935–2943 (2007).

7 Walsh, B. S. et al. The impact of climate change on fertility. Trends in Ecology & Evolution 34, 249–259 (2019).

8 Inouye, D. W. Climate change and phenology. Wiley Interdiscip Rev Clim Change 13 (2022). 10.1002/wcc.764

9 Amasino, R. Vernalization, Competence, and the Epigenetic Memory of Winter. Plant Cell 16, 2553–2559 (2004). 10.1105/tpc.104.161070 PMID - 15466409

10 Kim, D.-H. Current understanding of flowering pathways in plants: focusing on the vernalization pathway in Arabidopsis and several vegetable crop plants. Hortic Environ Biotechnology 61, 209–227 (2020). 10.1007/s13580-019-00218-5

11 Rogers-Bennett, L., Dondanville, R. F., Moore, J. D. & Vilchis, L. I. Response of Red Abalone Reproduction to Warm Water, Starvation, and Disease Stressors: Implications of Ocean Warming. J Shellfish Res 29, 599–611 (2010). 10.2983/035.029.0308

12 Breckels, R. D. & Neff, B. D. The effects of elevated temperature on the sexual traits, immunology and survivorship of a tropical ectotherm. J Exp Biol 216, 2658–2664 (2013). 10.1242/jeb.084962 PMID - 23531818

13 Paxton, C. W., Baria, M. V. B., Weis, V. M. & Harii, S. Effect of elevated temperature on fecundity and reproductive timing in the coral Acropora digitifera. Zygote 24, 511–516 (2016). 10.1017/s0967199415000477 PMID - 26349407

14 Hurley, L. L., McDiarmid, C. S., Friesen, C. R., Griffith, S. C. & Rowe, M. Experimental heatwaves negatively impact sperm quality in the zebra finch. Proceedings of the Royal Society B: Biological Sciences 285, 20172547 (2018). 10.1098/rspb.2017.2547 PMID - 29343605

15 Parratt, S. R. et al. Temperatures that sterilize males better match global species distributions than lethal temperatures. Nat Clim Change 11, 481–484 (2021). 10.1038/s41558-021-01047-0

16 Okamoto, D. K., Schmitt, R. J., Holbrook, S. J. & Reed, D. C. Fluctuations in food supply drive recruitment variation in a marine fish. Proceedings of the Royal Society B: Biological Sciences 279, 4542–4550 (2012). 10.1098/rspb.2012.1862 PMID - 23015631

17 Stoner, A. W. & Ray-Culp, M. Evidence for Allee effects in an over-harvested marine gastropod: density-dependent mating and egg production. Mar Ecol Prog Ser 202, 297–302 (2000). 10.3354/meps202297

18 Gaines, S. D. & Bertness, M. D. Dispersal of juveniles and variable recruitment in sessile marine species. Nature 360, 579–580 (1992). 10.1038/360579a0

19 Kelly, M. S. Environmental parameters controlling gametogenesis in the echinoid Psammechinus miliaris. Journal of Experimental Marine Biology and Ecology 266, 67–80 (2001).

20 Rogers-Bennett, L., Klamt, R. & Catton, C. A. Survivors of climate driven abalone mass mortality exhibit declines in health and reproduction following kelp forest collapse. Frontiers in Marine Science 8, 725134 (2021).

21 Okamoto, D. K., Schroeter, S. C. & Reed, D. C. Effects of ocean climate on spatiotemporal variation in sea urchin settlement and recruitment. Limnol Oceanogr 65, 2076–2091 (2020). 10.1002/lno.11440

22 Ebert, T. A., Schroeter, S. C., Dixon, J. D. & Kalvass, P. Settlement patterns of red and purple sea urchins (Strongylocentrotus franciscanus and S. purpuratus) in California, USA. Mar Ecol Prog Ser, 41–52 (1994).

23 Farmanfarmaian, A. & Giese, A. C. Thermal tolerance and acclimation in the western purple sea urchin, Strongylocentrotus purpuratus. Physiological Zoology 36, 237–243 (1963).

24 Azad, A. K., Pearce, C. M. & McKinley, R. S. Influence of stocking density and temperature on early development and survival of the purple sea urchin, Strongylocentrotus purpuratus (S timpson, 1857). Aquaculture Research 43, 1577–1591 (2012).

25 Azad, A. K., Pearce, C. M. & McKinley, R. S. Influence of microalgal species and dietary rations on larval development and survival of the purple sea urchin, Strongylocentrotus purpuratus (Stimpson, 1857). Aquaculture 322, 210–217 (2011).

26 Strathmann, M. F. Reproduction and development of marine invertebrates of the northern Pacific coast: data and methods for the study of eggs, embryos, and larvae. (University of Washington Press, 2017).

27 Basch, L. V. & Tegner, M. J. Reproductive responses of purple sea urchin (Strongylocentrotus purpuratus) populations to environmental conditions across a coastal depth gradient. Bulletin of Marine Science 81, 255–282 (2007).

28 Pearse, J. S. Synchronization of gametogenesis in the sea urchins Strongylocentrotus purpuratus and S Franciscanus in Advances in Invertebrate Reproduction (ed S. Adams W. H. Clark Jr) (Elsevier, 1981).

29 Hobday, A. J. et al. A hierarchical approach to defining marine heatwaves. Progress in Oceanography 141, 227–238 (2016).

30 Gronau, Q. F., Singmann, H. & Wagenmakers, E.-J. bridgesampling: An R package for estimating normalizing constants. Journal of Statistical Software 92, 1–29 (2020). doi:10.18637/jss.v092.i10

31 Byrne, M. Annual reproductive cycles of the commercial sea urchin Paracentrotus lividus from an exposed intertidal and a sheltered subtidal habitat on the west coast of Ireland. Marine Biology 104, 275–289 (1990).

32 Huey, R. B. & Kingsolver, J. G. Climate warming, resource availability, and the metabolic meltdown of ectotherms. The American Naturalist 194, E140–E150 (2019).

33 Cochran, R. C. & Engelmann, F. Environmental regulation of the annual reproductive season of Strongylocentrotus purpuratus (Stimpson). The Biological Bulletin 148, 393–401 (1975).

34 Spindel, N. B. Ecophysiology of Ectothermic Ecosystem Engineers: Bioenergetic Effects of Climate and Food on Dominant Consumers and Their Consequences for Coastal Ecosystems PhD thesis, The Florida State University, (2023).

35 Spindel, N. B. Ecophysiology of ectothermic ecosystem engineers: bioenergetic effects of climate and food on dominant consumers and their consequences for coastal ecosystems PhD thesis, Florida State University, (2023).

36 Cushing, D. Plankton production and year-class strength in fish populations: an update of the match/mismatch hypothesis in Advances in marine biology Vol. 26 249–293 (Elsevier, 1990).

37 Kapsenberg, L., Okamoto, D. K., Dutton, J. M. & Hofmann, G. E. Sensitivity of sea urchin fertilization to pH varies across a natural pH mosaic. Ecology and Evolution 7, 1737–1750 (2017).

38 Tegner, M. & Dayton, P. Sea urchins, E. Niños, and the long term stability of Southern California kelp forest communities. Marine Ecology Progress Series. 77, 49–63 (1991).

39 Zhadan, P. M., Vaschenko, M. A. & Almyashova, T. N. Spawning failure in the sea urchin Strongylocentrotus intermedius in the northwestern Sea of Japan: Potential environmental causes. Journal of Experimental Marine Biology and Ecology 465, 11–23 (2015).

40 Seward, L. C. The relationship between green sea urchin spawning, spring phytoplankton blooms, and the winter-spring hydrography at selected sites in Maine MSc thesis, University of Maine, (2002).

41 Lester, S. E., Tobin, E. D. & Behrens, M. D. Disease dynamics and the potential role of thermal stress in the sea urchin, Strongylocentrotus purpuratus. Canadian Journal of Fisheries and Aquatic Sciences 64, 314–323 (2007).

42 Behrens, M. D. & Lafferty, K. D. Effects of marine reserves and urchin disease on southern Californian rocky reef communities. Mar Ecol Prog Ser 279, 129–139 (2004).

43 Lafferty, K. D. Fishing for lobsters indirectly increases epidemics in sea urchins. Ecological Applications 14, 1566–1573 (2004).

44 Arafeh-Dalmau, N. et al. Extreme marine heatwaves alter kelp forest community near its equatorward distribution limit. Frontiers in Marine Science 6, 499 (2019).

45 Filbee-Dexter, K. et al. Marine heatwaves and the collapse of marginal North Atlantic kelp forests. Scientific Reports 10, 13388 (2020).

46 McPherson, M. L. et al. Large-scale shift in the structure of a kelp forest ecosystem co-occurs with an epizootic and marine heatwave. Communications Biology 4, 298 (2021).

47 Michaud, K. M., Reed, D. C. & Miller, R. J. The Blob marine heatwave transforms California kelp forest ecosystems. Communications Biology 5, 1143 (2022).

48 Smale, D. A. Impacts of ocean warming on kelp forest ecosystems. New Phytologist 225, 1447–1454 (2020).

49 Thomsen, M. S. et al. Local extinction of bull kelp (Durvillaea spp.) due to a marine heatwave. Frontiers in Marine Science 6, 84 (2019).

50 Wernberg, T. Marine heatwave drives collapse of kelp forests in Western Australia in Ecosystem Collapse and Climate Change (eds J. G. Canadell & R. B. Jackson) 325–343 (Springer, 2021).

51 Smith, K. E. et al. Socioeconomic impacts of marine heatwaves: Global issues and opportunities. Science 374, eabj3593 (2021).

52 Walsh, B. S. et al. Female fruit flies cannot protect stored sperm from high temperature damage. J Therm Biol 105, 103209 (2022).

53 Carter, M. L. et al. Shore Stations Program, La Jolla - Scripps Pier (La Jolla Archive, 2025-03-14) in Shore Stations Program Data Archive: Current and Historical Coastal Ocean Temperature and Salinity Measurements from California Stations. UC San Diego Library Digital Collections. (2022).

54 Carter, M. L. et al. (UC San Diego Library Digital Collections, 2022).

55 Reynolds, R. W. et al. Daily High-Resolution-Blended Analyses for Sea Surface Temperature. J Climate 20, 5473–5496 (2007). 10.1175/2007jcli1824.1

56 Kass, R. E. & Raftery, A. E. Bayes factors. Journal of the American Statistical Association 90, 773–795 (1995).

57 Meng, X.-L. & Wong, W. H. Simulating ratios of normalizing constants via a simple identity: a theoretical exploration. Statistica Sinica, 831–860 (1996).

58 Stan User Guide v. 2.36 (2025).

59 Bürkner, P.-C. brms: An R package for Bayesian multilevel models using Stan. Journal of Statistical Software 80, 1–28 (2017).

